# Incomplete transcriptional dosage compensation of vertebrate sex chromosomes is balanced by post-transcriptional compensation

**DOI:** 10.1101/2023.02.23.529605

**Authors:** Nicholas C Lister, Ashley M Milton, Hardip R Patel, Shafagh A Waters, Benjamin J Hanrahan, Kim L McIntyre, Alexandra M Livernois, Lee Kian Wee, Alessa R. Ringel, Stefan Mundlos, Michael I. Robson, Linda Shearwin-Whyatt, Frank Grützner, Jennifer A. Marshall Graves, Aurora Ruiz-Herrera, Paul D Waters

## Abstract

Heteromorphic sex chromosomes (XY or ZW) present problems of gene dosage imbalance between the sexes, and with the autosomes. Mammalian X chromosome inactivation was long thought to imply a critical need for dosage compensation in vertebrates. However, the universal importance of sex chromosome dosage compensation was questioned by mRNA abundance measurements that demonstrated sex chromosome transcripts are neither balanced between the sexes or with autosomes in monotreme mammals or birds. Here, we demonstrate unbalanced mRNA levels of X genes in platypus males and females that correlate with differential loading of histone modifications, and confirm that transcripts of Z genes are unbalanced between males and females also in chicken. However, we found that in both species, median male to female protein abundance ratios were 1:1, implying an additional level of post-transcriptional control. We conclude that parity of sex chromosome output is achieved in birds, as well as all mammal groups, by a combination of transcriptional and post-transcriptional control, consistent with an essential role for sex chromosome dosage compensation in vertebrates.

## Introduction

Therian (eutherian and marsupial) mammals have XY males and XX females in which X-borne genes are unequally represented between the sexes. Birds have ZZ males and ZW females in which Z-borne genes are unequally represented. Thus, despite their different sex chromosome systems, mammals and birds (and many other vertebrates and invertebrates with differentiated sex chromosomes) share the challenge that dosage of sex-linked genes is unequal between males and females, and between sex chromosomes and autosomes.

Platypus (an egg-laying monotreme mammal) has an even more acute dosage problem arising from a complex sex chromosome system, in which males have five X chromosomes and five degraded Y chromosomes, and females have five pairs of X chromosomes^1^. These sex chromosomes form a chain at meiosis held together by recombination within nine pseudoautosomal regions (PARs) shared between successive X and Y chromosomes. The platypus sex chromosomes have no homology with the sex chromosomes of therian mammals, but share considerable homology with the chicken Z^2^.

In therian mammals, parity of expression of X-borne genes between the sexes is achieved by the epigenetic silencing of one X in the somatic cells of females. X chromosome inactivation (XCI) was long assumed to be at the transcriptional level, and this was experimentally demonstrated^3^ and repeatedly confirmed^4,5^. XCI is considered to be a global silencing mechanism, although many genes escape inactivation on the human X.

Unequal expression of X-borne and autosomal genes in males has been suggested to be compensated by upregulation of X-borne genes, a step considered by Ohno^6^ to have driven the evolution of XCI. However, reports of X upregulation in human and mouse are disputed^4,7-9^, and it may be that only a subset of some dosage-sensitive X-borne genes are upregulated^7^. Global upregulation of the active X occurs in only a few species^4,7^. In eutherian mammals, X to autosome transcriptional output is less than 1 in both sexes^4^ but this reduced X expression relative to the autosomes is compensated in the proteome, at least in males^10^.

Understanding of the complex and stable XCI system in therian mammals promoted the view that dosage compensation between sexes is essential for any animal with differentiated sex chromosomes. However, this view was challenged by studies of platypus and birds. In platypus, male:female (M:F) ratios of X-borne transcripts fell between complete compensation (1.0) and complete absence of dosage compensation (0.5), with a median of 0.6^4^. Similarly in birds, transcription from Z-borne genes showed M:F ratios between full compensation (1.0) and absence of compensation (2.0), with a median of 1.4^4,11^. Evidently, Z chromosome-wide dosage compensation does not occur in birds, and might be restricted to dosage sensitive genes^4,12^.

The absence of chromosome wide compensation at the transcriptional level in monotremes and birds provoked much discussion about the universal importance of dosage compensation^7,13,14^. Likewise, there is conflicting evidence of sex chromosome/autosome dosage compensation at the level of the proteome (i.e., protein abundance)^7,15,16^. Here, we investigated dosage compensation in both the transcriptome and proteome of representative therian mammals; (mouse and opossum), monotremes (platypus) and birds (chicken). We demonstrate that incomplete transcriptional dosage compensation of sex chromosomes is balanced by post-transcriptional compensation in platypus and chicken.

### Compensation by transcriptional and post-transcriptional control

We first measured global mRNA and protein output from X-borne genes in mouse and opossum and calculated the M:F ratios of RNA and protein abundance. We found that transcriptional output was equivalent (M:F ∼ 1) between the sexes in mouse and opossum (Extended Data Fig 1), as previously described^4^, and a 1:1 ratio was also observed in their respective proteomes. Thus, male to female X chromosome gene dosage equivalence is established in the transcriptome and parity is maintained in the proteome in therian mammals.

However, this was not the case for platypus sex chromosomes. The need for dosage compensation in the platypus would be expected to be acute, given that 9.23% of the genome (the non-PARs of the five X chromosomes, Fig 1) is present in a single dose in males. We defined seven of the nine PARs to 100kbp resolution using male fibroblast HiC interaction patterns (Fig 1), with X_3_Y_2_ defined to 500kbp resolution. The PARs are all assembled on X chromosome contigs, with Y contigs bearing only male specific sequence. Because Y-specific sequences are adjacent to their associated PARs, genomic interactions with the Xs clearly demarcated PAR boundaries for all except the tiny X_4_Y_4_ PAR.

**Fig 1.**
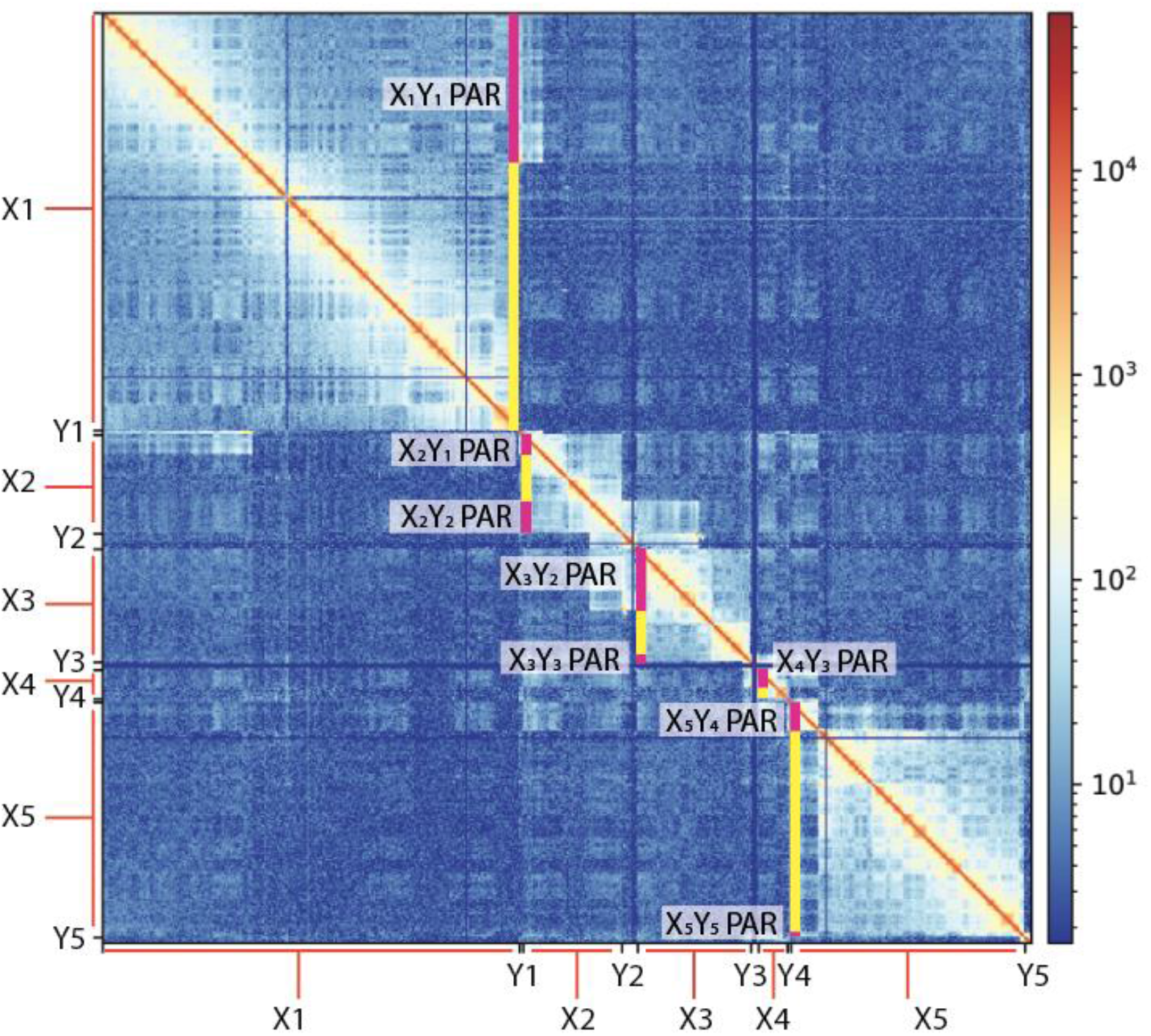
HiC data of the platypus X chromosomes assembled to define PAR boundaries. The PARs in platypus are assembled on the X chromosome contigs. Y contigs in the assembly only contain Y specific sequence. Hi-C interactions between the PARs on the Xs and Y specific material were used to define PAR boundaries, with proximal PARs having increased contacts with their corresponding Ys compared to distal PAR regions. The HiC contact matrix for X and Y chromosomes in male platypus were generated using 100 kb bins. The heatmap scale indicates the number of contacts between bins. PARs are shown with purple bars and X specific regions by yellow bars.

We measured RNA and protein output from autosomes and X chromosomes in platypus fibroblasts, heart and liver, differentiating between X-specific regions and PARs. M:F expression ratios of RNA and protein were close to 1:1 for genes on autosomes and in the PARs of X_1_ and X_5_. However, transcription of genes on the X-specific portion of X_1_ was significantly lower in males than females, with a median M:F ratio of 0.7-0.75 across the three tissue types (Fig 2A). Likewise, the X-specific region of X_5_ showed strong female bias. We observed the same pattern when all X specific, and all PAR genes were binned together (Fig 2A).

**Fig 2.**
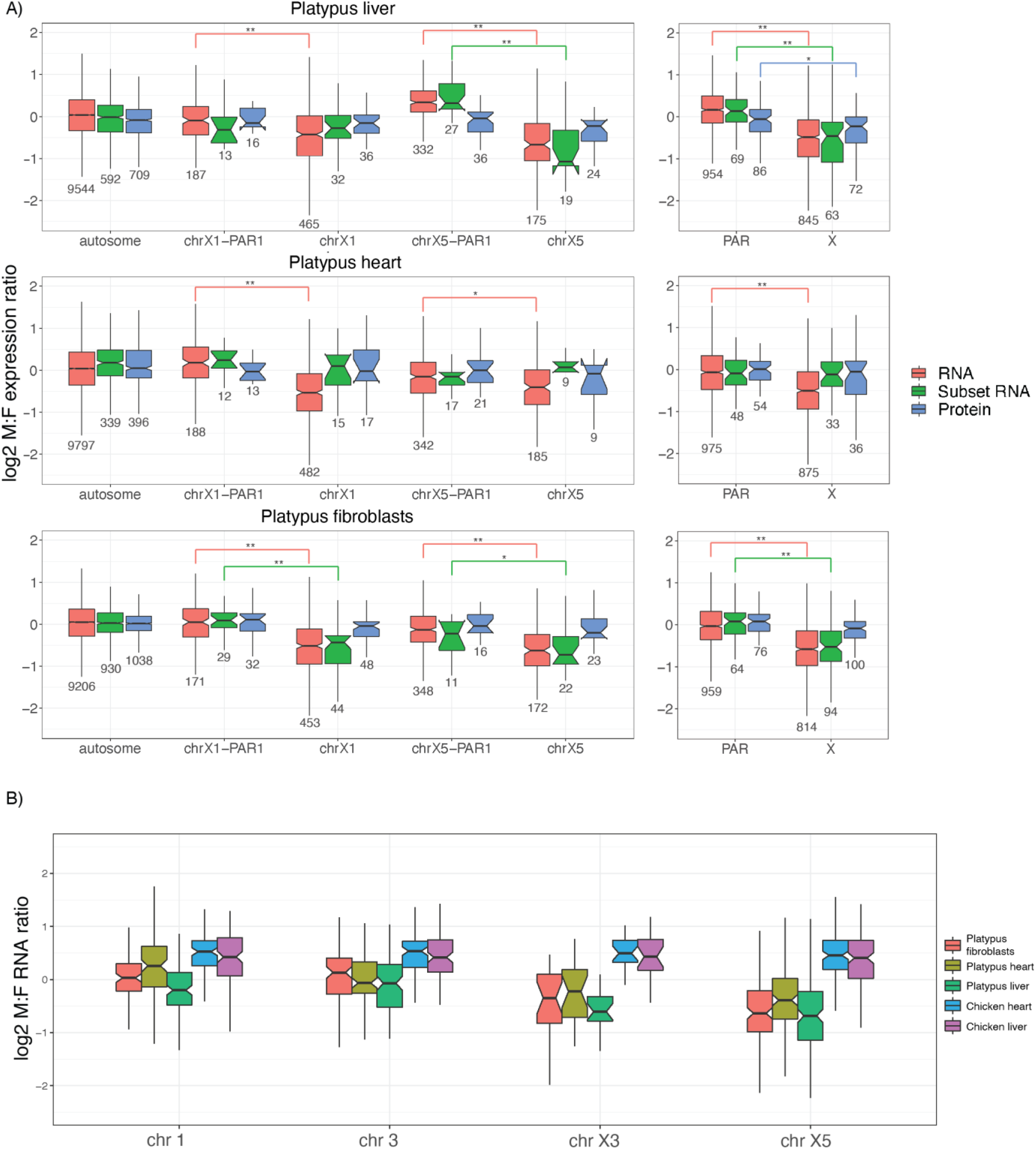
Male to female expression ratios of X borne and autosomal genes in platypus. Median male to female (M:F) transcript abundance ratios on a log2 scale of **A**) autosomal and X borne genes in platypus fibroblasts, heart and liver. On the far right, ratios are shown for all X specific genes and PAR genes. Ratios were calculated for all expressed mRNA (red), the subset of mRNAs sampled in the proteome (green), and proteins (blue). A ratio above zero is higher expression in males, whereas below zero is lower expression. Boxes represent the middle 50% of the data, and whiskers represent 1.5 times the interquartile range. Outliers are not plotted. Median is plotted inside the box, with the number of genes sampled below each boxplot. A Mood’s median test was used to calculate statistical difference (** p < 0.001, * p < 0.01). **B)** Male to female ratios in the transcriptome of orthologues sampled on the chicken Z and platypus. In chicken the Z genes have higher transcriptional output in males. Genes that are autosomal in platypus have equal expression between the sexes. Genes that are on a platypus X have higher median expression in females.

We then compared relative protein abundance of genes on the autosomes, the PARs and X-specific regions of X_1_ and X_5_ in platypus males and females. Genes in autosome and PAR regions showed a 1:1 ratio. Unexpectedly, we found that the M:F protein ratios were also 1:1 in both X specific regions, and in all X specific regions combined (Fig 2A). This was not due to sampling of a subset of mRNAs that were represented in the proteome (green boxplots in Fig 2A), since this subset had the same M:F ratios as the total transcriptome (red boxplots in Fig 2A). This implies that, as well as partial transcriptional regulation, dosage compensation in platypus occurs post-transcriptionally as well as transcriptionally.

To investigate if a similar pattern of DC occurs in birds, we profiled the transcriptome and proteome of chicken heart and liver. This showed that autosomal genes had M:F transcriptome ratios near 1:1, whereas Z-borne genes had a strong male bias, as previously reported^17,18^. However, the median M:F abundance ratios of proteins encoded by Z genes were near 1:1 in both tissues (Fig 2B, Extended Data Fig 2), implying that protein levels are fully compensated for all Z-borne genes, rather than restricted to 30% as previously reported^16^.

These results demonstrate lack of global dosage compensation at the transcriptional level, but full dosage compensation at the protein level in platypus and chicken. This contrasts with therian mammals, where dosage compensation occurs solely in the transcriptome.

### Molecular basis of partial transcriptional dosage compensation in platypus

XCI in eutherians involves DNA methylation and histone modifications^19,20^. We analysed published DNA methylation data for platypus^21^, finding no difference in DNA methylation levels between the sexes on the X chromosomes globally (Extended Data Fig 3).

We also performed Chromatin Immunoprecipitation (ChIP) in platypus fibroblasts to detect histone modifications that are associated with active transcription: H3K27ac, H3K4me1, H3K4me3 and H4K20me1. In both sexes, fewer ChIP-peaks were detected in X specific regions compared to autosomes and PARs for all four histone modifications marks (Fig 3, Extended Data Fig 4, Extended Data Fig 5, Table 1). For H3K27ac, H3K4me3 and H4K20me1, genome wide ChIP-peak density (peaks per Mbp) was comparable between sexes (Table 1). However, for H3K4me1 the X chromosomes showed a striking difference between sexes, an 89% reduction in X to autosome peak density in females compared to only a 19% reduction in males (Table 1). The higher density of this active histone mark in male platypus could activate transcription of the Xs. Thus, partial transcriptional upregulation of X chromosomes in males correlates with differential loading of the active histone modification H3K4me1 between sexes.

**Table 1.** ChIP peak density.

| Modification | Peaks/Mbp |  |  |  |  |  |
| --- | --- | --- | --- | --- | --- | --- |
|  | Male |  | Male A/X | Female |  | Female A/X |
|  | Autosome | Male X | ratio | Autosome | X | ratio |
| H3K4me1 | 26.25 | 21.34 | 0.81 | 44.58 | 5.12 | 0.11 |
| H3K4me3 | 21.42 | 10.6 | 0.49 | 24.76 | 7.88 | 0.32 |
| H3K27ac | 21.37 | 5.46 | 0.26 | 30.91 | 13.82 | 0.45 |
| H4K20me1 | 28.96 | 9.69 | 0.33 | 24.5 | 14.22 | 0.58 |

**Fig 3.**
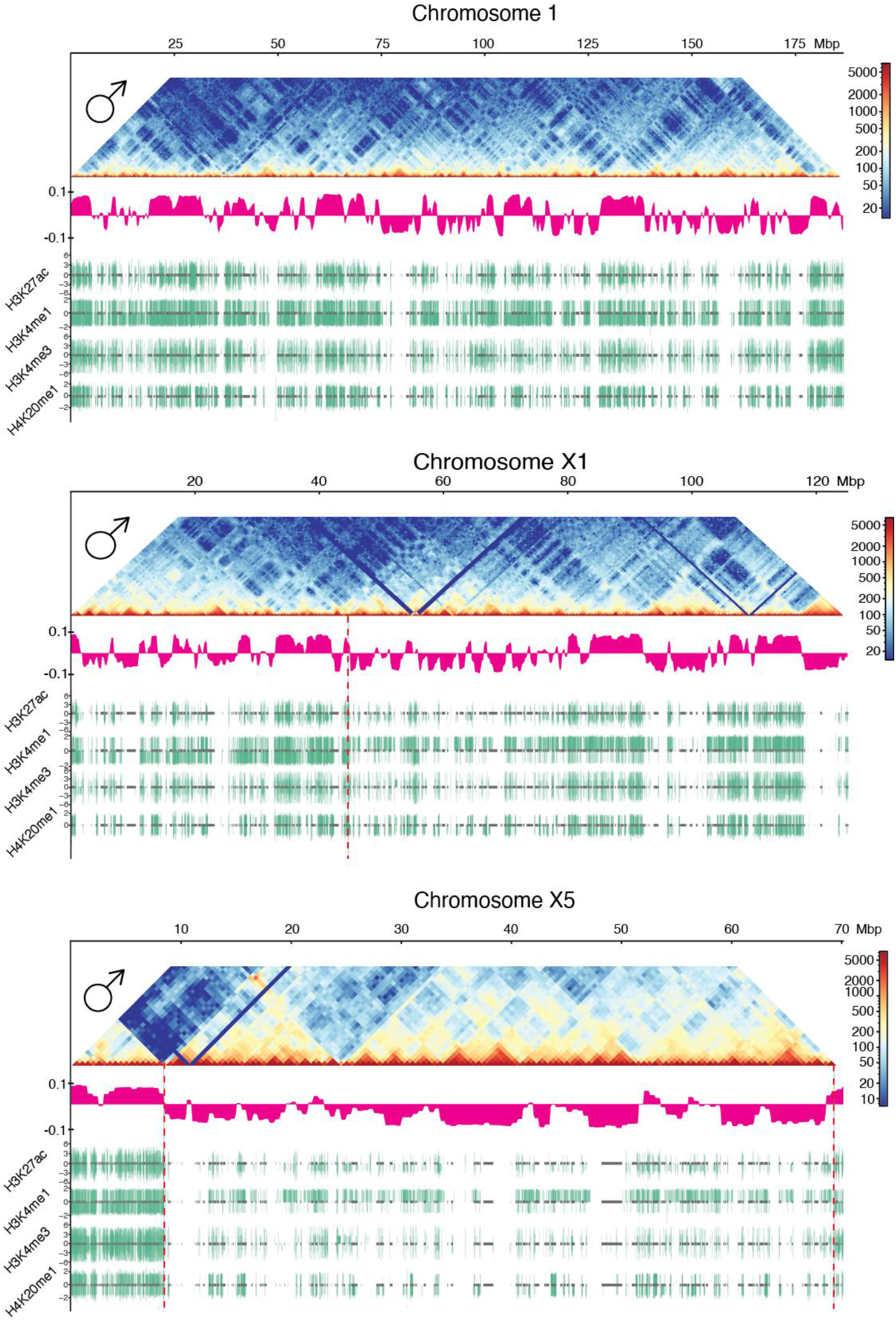
Epigenetic profile and 3D structure of platypus chromosomes 1, X1 and X5. Whole chromosome plots for chromosome 1 (a representative autosome), X1 and X5 in platypus. The x-axis represents the full length of the chromosome, with size (in Mbp) at the top. Each plot shows a Hi-C contact matrix heatmap for male (top) generated using 500 kb bins. The Hi-C heatmap scale indicates number of contacts between bins. Immediately below the Hi-C heatmaps are eigenvectors (pink tracks), which indicate A (positive) and B (negative) compartments. Below the eigenvectors are ChIP-seq peak tracks for four active histone marks (green): H3K27ac, H3K4me1, H3K4me3 and H4K20me1. For each histone mark, male and female peaks are shown above and below the centre line respectively. ChIP peaks were filtered using a threshold q-value of 0.05 and peak height is displayed up to a 5-fold change. Gene locations are shown in the centre of each ChIP track as grey boxes. Lines are the median M:F ratios for the whole chromosome. On the X chromosome plots, vertical dashed red lines indicate PAR boundaries that were identified with HiC interaction data (see Fig 1).

### Full compensation by control at multiple regulatory levels

Here we demonstrate that sex chromosome gene dosage is compensated fully (or nearly so) by transcriptional control in therian mammals, but not in platypus or chicken (Fig 4). Our most significant finding is that although X and Z borne genes in platypus and chicken are variably and only partially compensated by differential transcription, they are fully compensated between the sexes at the protein level.

**Fig 4.**
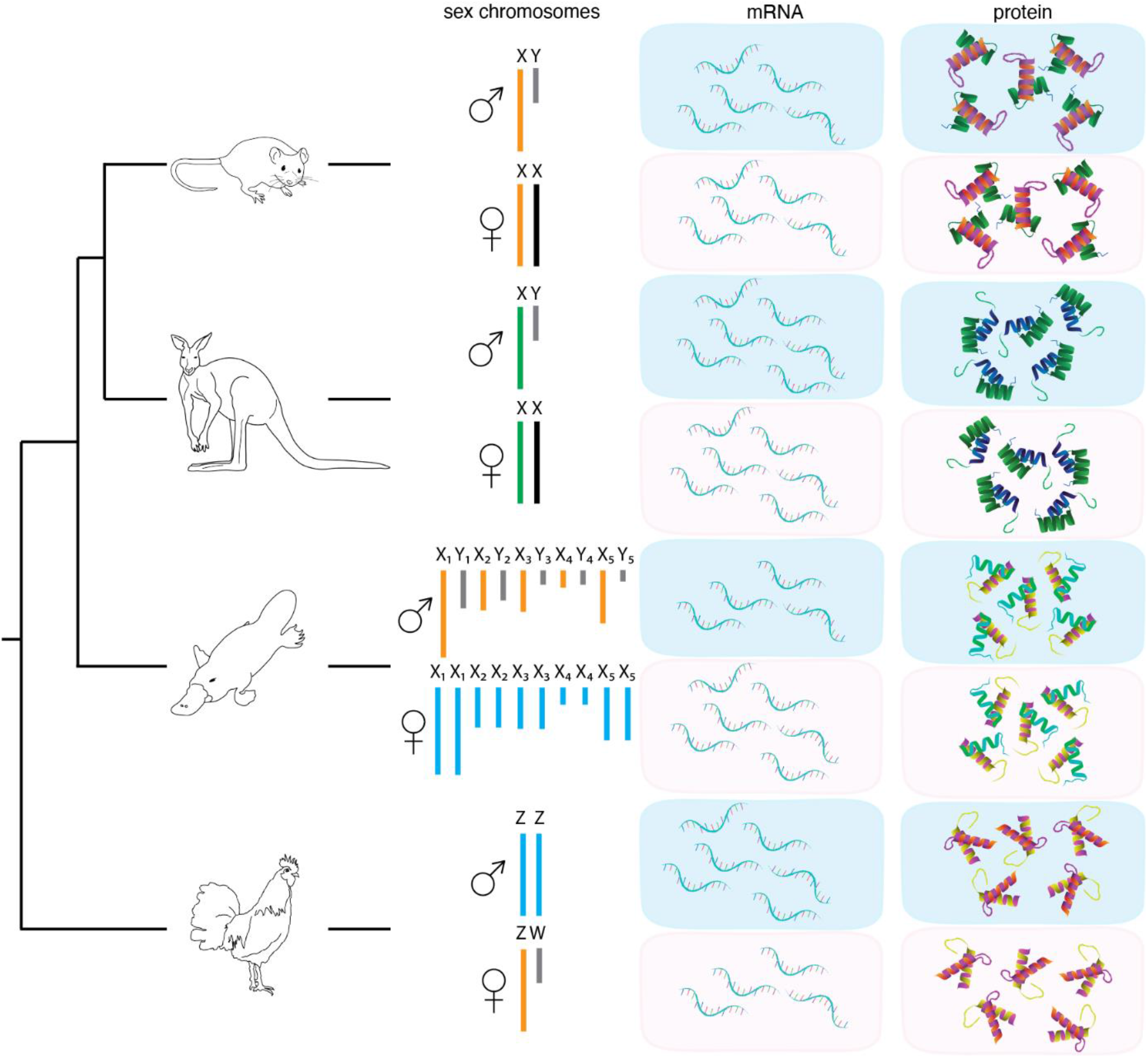
Transcriptome and proteome sex chromosome dosage compensation. Relationships of the different study species and their respective sex chromosome and dosage compensation systems. For each species, equal or unequal output between the sexes is indicated by the number of molecules shown for the transcriptome (mRNA) and proteome (protein). The green X in marsupials is upregulated to match autosome transcriptional output. The orange X/Z in mouse, platypus and chicken indicates partial upregulation, while blue X/Z in platypus and chicken indicates no known partial upregulation in the homogametic sex. Grey chromosomes represent the degraded sex-specific Ys or Ws. Black chromosomes indicate X/Z has undergone X chromosome inactivation

There are two important implications of our findings. Firstly, genes sampled in the proteome must be up-regulated post-transcriptionally from the X/Z in the heterogametic sex, or downregulated in the homogametic sex, either by differential translation or altered transcript decay rates of X/Z genes in males and females and relative to autosomal genes^22^. Secondly, this regulation must be gene-specific because different genes are transcriptionally regulated to different extents.

We know of no established mechanism by which transcripts can be recognised by the translation mechanism as originating from sex chromosomes. Recognition could involve epigenetic modifications of RNA^23^ or sex biased microRNAs (miRNA), which have the potential to down regulate the expression of many genes. Sex-biased miRNAs have a high turnover rate, and examples were found in birds for miRNAs that have copies present on both the Z and W, and for miRNAs that preferentially down-regulate Z genes in males^24^.

Thus, far from dosage compensation being optional in non-therian vertebrates, our results suggest it is essential. However, it is accomplished by a combination of transcriptional and post-transcriptional control, rather than the transcriptional repression of one X chromosome in therian mammals.

### Evolution of dosage compensation in vertebrates

Is this combination of transcriptional and post-transcriptional control ancestral to vertebrates, or at least to amniotes? Transcriptional inactivation, although common to all three mammal branches, shows differences at the molecular level. XCI in eutherian mammals involves several histone modifications as well as DNA methylation, coordinated by an inactivation centre that bears multiple genes transcribed into noncoding RNAs, whereas transcriptional X inactivation in marsupials uses a different but overlapping set of histone modifications^25^, a different DNA methylation profile^21^ and an unrelated noncoding RNA^26^. In platypus, partial transcriptional upregulation was correlated to sex differences in one of the four active histone marks, suggesting a different use of ancient epigenetic mechanisms. Post-transcriptional control of gene activity complements transcriptional control only in non-therian vertebrates, but may play a role in regulation of the autosome-sex chromosome balance in eutherians^10^.

Thus our observations reveal different evolutionary strategies to compensate for different sex chromosome gene dosage between males and females in different vertebrate lineages (Figure 4). Evolutionary flexibility can be seen by comparing expression of orthologous genes on the chicken Z and platypus X chromosomes, which have higher expression in ZZ male birds, but lower expression in XY male platypuses (Fig 2B).

It is interesting to consider how these different modes of dosage compensation evolved in mammals. The balancing of incomplete transcriptional dose regulation by post-transcriptional control in birds as well as monotremes suggests that both mechanisms were involved in dosage compensation in a common reptilian ancestor. It will be instructive to compare dosage compensation in non-avian reptiles, amphibians and fish. The Komodo dragon has a ZW system with no dosage compensation of Z genes^27^. Conversely, the green anole has an XY system with complete dosage compensation of genes in the most differentiated region of the X, and incomplete dosage compensation of genes on newer, less differentiated regions of the X^28^. The dominance of transcriptional repression of X-borne genes, specific to therian mammals, would appear to be a later evolutionary event, perhaps relying on sequences specific to a relatively young therian X chromosome.

The different dosage compensation mechanisms of vertebrate groups may therefore represent different mixes of ancient epigenetic mechanisms that affect transcriptional and post-transcriptional control of gene activity.

## Methods

### Lead Contact

Further information and requests for resources and reagents should be directed to and will be fulfilled by the Lead Contact, Paul Waters.

### Materials Availability

This study did not generate new unique reagents.

## EXPERIMENTAL MODEL AND SUBJECT DETAILS

### Ethical Guidelines

All animal experiments were approved by the Australian National University Animal Experimentation Ethics Committee (approval number R.CG.14.08) and the Garvan/St Vincents Animal Ethics Committee (#13/35).

### Chicken Heart and Liver Tissue

For both male and female chicken (*Gallus gallus*), 3 biological replicates were used for heart tissues while 2 biological replicates were used for liver.

### Mouse Heart and Liver Tissue

A total of 2 biological replicates were used for both heart and liver samples in male and female mouse (*Mus musculus*).

### Opossum and Platypus Fibroblasts

Primary fibroblast cell cultures were established from ear clips of male and female individuals of both opossum (*Monodelphis domestica*) and platypus (*Ornithorhynchus anatinus*). Opossum fibroblast cell lines were cultured at 35°C and platypus fibroblasts at 37°C in 5% CO_2_ with DMEM containing 10% *(v/v)* FBS and 10ml/L PSG.

## METHOD DETAILS

### RNA extraction

One million fibroblast cells and 25mg of tissue from either heart or liver was used for RNA isolation using TRIzol reagent (ThermoFisher Scientific) and purified using RNeasy kit (Qiagen).

### Protein extraction

Proteins were extracted from heart and liver tissues of chicken, mouse and platypus, as well as fibroblasts for platypus and opossum, using RIPA lysis buffer (150mM NaCl, 1% NP-40, 0.5% sodium deoxycholate, 0.1% SDS, 2mM EDTA, 50mM Tris HCl, pH 8.0). The tissues were homogenized to encourage lysis of cells. The lysates were placed on ice for 30 minutes before being centrifuged at 10,000 x *g* for 20 minutes. The residual insoluble fraction of the extraction was subsequently extracted and analyzed on a 12% SDS-PAGE gel. Qubit quantification was performed to determine protein concentration. The mass spectrometry proteomics data have been deposited to the ProteomeXchange Consortium via the PRIDE^1^ partner repository with the dataset identifier PXD040182.

### LC-MS/MS

Chicken, mouse and platypus heart and liver samples were sent to Bioanalytical Mass Spectrometry Facility (BMSF) at the Mark Wainwright Analytical Centre (MWAC) at UNSW, while opossum and platypus fibroblast samples were sent to the Australian Proteome Analysis Facility (APAF) at Macquarie University.

Samples (dried) were solubilised in 1mL of 0.25M TEAB 0.05% SDS and buffer exchanged, using a Viva2 spin column (5kDa), into 250uL of 0.25M TEAB 0.05% SDS. Samples were quantified using Direct Detect (Millipore). 100ug of each sample (in duplicate) was reduced with TCEP, alkylated with MMTS and digested with trypsin according to SOP MS-001. The digested samples were labelled, cleaned and fractionated by SCX HPLC. The buffer A was 5mM Phosphate 25% Acetonitrile, pH 2.7 and buffer B was 5mM Phosphate 350mM KCL 25% Acetonitrile, pH 2.7. The dried iTRAQ labelled sample was resuspended in loading buffer which was the same as the buffer A. After sample loading and washing with buffer A, buffer B concentration increased from 10% to 45% in 70 minutes and then increased quickly to 100% and stayed at 100% for 10 minutes at a flow rate of 300 ul/min. The eluent of SCX was collected every 2 minutes at the beginning of the gradient and at 4 minutes interval later.

The labelled sample was resuspended in 100 µL of loading/desalting solution (0.1% formic acid and 2% acetonitrile 97.9% water). Sample (40µL) was injected onto a peptide trap (Michrome peptide Captrap) for pre-concentration and desalted with 0.1% formic acid, 2% ACN, at 5 µL/min for 10 minutes. The peptide trap was then switched into line with the analytical column. Peptides were eluted from the column using a linear solvent gradient, with steps, from mobile phase A: mobile phase B (98:2) to mobile phase A:mobile phase B (65:35) where mobile phase A is 0.1% formic acid and mobile phase B is 90% ACN/0.1% formic acid at 600 nL/min over a 100 min period. After peptide elution, the column was cleaned with 95% buffer B for 15 minutes and then equilibrated with buffer A for 25 minutes before next sample injection. The reverse phase nanoLC eluent was subject to positive ion nanoflow electrospray analysis in an information dependant acquisition mode (IDA). Page 3 of 4 Commercial in confidence In IDA mode a TOFMS survey scan was acquired (m/z 400 - 1500, 0.25 second), with the ten most intense multiply charged ions (counts >150) in the survey scan sequentially subjected to MS/MS analysis. MS/MS spectra were accumulated for 200 milliseconds in the mass range m/z 100 – 1500 with the total cycle time 2.3 seconds.

### ChIP-Seq

Platypus fibroblast cells were harvested and fixed with 1% formaldehyde in 10% FCS/PBS. for 10 min at RT. Crosslinking was quenched by adding glycine to a final concentration of 125 mM. To isolate nuclei the fixed cells were washed twice with cold PBS and lysed using fresh lysis buffer (10 mM Tris, pH 7.5, 10 mM NaCl, 5 mM MgCl2, 0.1 mM EGTA with protease inhibitor). Nuclei were centrifuged for 5 min /480g /4°C, washed with PBS and either snap frozen in liquid N_2_ or further processed. Chromatin for ChIP-seq was sheared using a Bioruptor until DNA fragment size of 200–500 base pairs was reached. Afterwards, samples were processed with the iDeal ChIP-seq kit for histones/transcription factor according to the manufacturer’s instructions. For each Histone ChIP 5 μg chromatin was used in combination with antibodies against H3K4me1 (1 µg) H3K4me3 (1 µg), H3K27ac (1 µg), and H4K20me1 (5 µg). For H3K4me1, K3K4me3 and H3K27ac, libraries were prepared for sequencing using the KAPA HyperPrep kit and their quality confirmed by Bioanalyzer analysis. Samples were run on NovaSeq 6000 platform.

For H4K20me1, 20µl of magnetic G beads (Millipore) were added to each ChIP sample and incubated at 4°C rotating for 2 hours. Samples were centrifuged briefly, and then beads were allowed to settle in a magnetic rack before removing supernatant. Beads were washed consecutively with 1ml of low salt buffer, 1ml of high salt buffer, 1ml of LiCI buffer, 1ml of TE buffer, for 5 minutes each at 4°C. Between each wash samples were centrifuged briefly, and then beads were allowed to settle in a magnetic rack before removing supernatant. After the final wash, tubes were spun briefly and contents were transferred fresh 1.5ml tube. Beads were allowed to settle in a magnetic rack, and supernatant was removed. Tubes were removed from magnetic rack and 100µl of elution buffer was added, followed by agitation at room temperature for 30 minutes. Beads were allowed to settle in a magnetic rack and supernatant (chromatin-antibody complexes) was transferred to a fresh tube. NaCl was added to final concentration of 0.6M. Total input was thawed and diluted with 4 volumes elution buffer, and NaCl was added to a final concentration of 0.6M. Samples were incubated at 67°C overnight to reverse cross-links. Samples were sent to the Beijing Genomics Institute (BGI) for library construction and sequencing on HiSeq 2000 platform.

### Hi-C

Hi-C libraries were prepared as described in a previously published *in situ* protocol (Melo et al., 2020; Rao et al., 2014). Briefly, male platypus fibroblast cells were harvested and fixed with 2% formaldehyde in 10% FCS/PBS for 10 min at RT. Crosslinking was quenched by adding glycine to a final concentration of 125 mM. To isolate nuclei the fixed cells were washed twice with cold PBS and lysed using fresh lysis buffer (10 mM Tris, pH 7.5, 10 mM NaCl, 5 mM MgCl2, 0.1 mM EGTA with protease inhibitor). Nuclei were centrifuged 5 min /500g / 4°C, washed several times with PBS and subsequently digested overnight with DpnII enzyme. Digested DNA ends were marked with biotin-14-dATP and ligated overnight using T4 DNA ligase. Formaldehyde crosslinking was reversed by incubation in 5 M NaCl for 2 h at 68°C, followed by ethanol precipitation. Chromatin was sheared into 300–600 bp fragments using a S-Series 220 Covaris for library preparation, afterwards biotin-filled DNA fragments were pulled down using Dynabeads MyOne Streptavidin T1 beads. DNA ends were repaired using T4 DNA polymerase and the Klenow fragment of DNA polymerase I and phosphorylated with T4 Polynucleotide Kinase NK. Afterwards adaptors were ligated to DNA fragments using the NEBNext Multiplex Oligos for Illumina kit. Indexes were added via PCR amplification (3–8 cycles) using the NEBNext Ultra II Q5 Master Mix. PCR purification and fragment size selection were done using Agencourt AMPure XP beads. Library sequencing was done on a NovaSeq 6000 platform.

## QUANTIFICATION AND STATISTICAL ANALYSIS

### Sequencing

Chicken and mouse samples were analysed by 1 × 75 bp single-read sequencing on Illumina NextSeq 500. Library construction and Illumina sequencing performed by Ramaciotti Centre for Genomics (UNSW Sydney, Australia), using the TruSeq Stranded mRNA Library Prep Kit. Opossum and platypus samples were analysed by 1x 100 bp paired end sequencing on Illumina HiSeq 2000 by Beijing Genomics Institute (BGI). The resulting raw sequence data is available on the NCBI short read archive as BioProject PRJNA929280.

### Sequence mapping

The raw reads from RNASeq, Hi-C and ChIP-seq were analysed using FastQC^2^ to assess for quality.

Reference genomes were masked for the Y chromosome sequences in platypus and mouse and for the W chromosome in chicken to reduce false positive mappings in the sex lacking these chromosomes for RNASeq data. This was not required for the opossum assembly which was from a female sample. The reference genome versions were GRCg6a for chicken, mOrnAna1.p.v5.1 for platypus, MonDom5 for opossum and GRCm39 for mouse. Subread^3^ was used to align the reads to their reference genome. For RNASeq reads, parameters were chosen such that (a) all possible subreads were chosen for mapping (-n 300) and (b) for multi-mapping reads, one random location was chosen for read assignments (--multiMapping -B 1). For Hi-C reads, Subread was run in single-end mode to generate separate forward and reverse .sam files, which were required in subsequent processing. All parameters were left as default. Subread was used on ChIP-seq reads in the same fashion, with the exception of specifying a Phred score of phred+64 (-P 6). To account for differences in number of mapped reads between male and female Hi-C samples, Samtools^4^ version 1.11 was used to take a subset of the male Hi-C reads. 1.7% of male reads were randomly chosen, to be equivalent to the number of female reads. Unless otherwise stated, this male subset was used in all subsequent analyses.

### Counting mapped reads

FeatureCounts^5^ was used to calculate the number of RNASeq reads assigned to each feature based on gtf files from NCBI. Parameters were chosen such that (a) reads were assigned to all overlapping features if features were overlapping (-O), (b) multi-mapping reads were counted (-M), and (c) strand-specificity of RNAseq was accounted for counting (-s 0,1, or 2). For paired-end data, fragments were counted instead of individual reads of a pair (-p).

### ProteinPilot software workflow

The MS/MS data were submitted to ProteinPilot V5.0.2 (AB Sciex)^6^ for data processing, using databases obtained from SwissProt. Databases relevant for each species were used. Bias correction was selected. The detected protein threshold (unused ProtScore) was set as larger than 1.3 (better than 95% confidence). The Protein Pilot group file and the protein summaries were exported and male:female ratios calculated.

In platypus, two heart samples each from male (113 and 114) and female (115 and 116), and two liver samples each from male (117 and 118) and female (119 and 121) were analyzed. In chicken, three heart samples each from male (113, 114 and 115) and female (116, 117 and 118), and two liver samples each from male (113 and 114) and female (115 and 116) were analyzed. In mouse, two heart and liver samples each from male (117 and 118) and female (119 and 121) were analyzed. One sample each from platypus fibroblasts were labelled twice for male (115 and 117) and female (114 and 116). One sample each from opossum fibroblasts were labelled twice for male (114 and 116) and female (115 and 117). Male:female ratios between each male and female sample were used to calculate geomeans for each gene.

### Hi-C matrix generation and analysis

Hicexplorer^7-9^ version 3.4.3 was used for Hi-C matrix generation and analysis. HicBuildMatrix was used to generate 500 kb .h5 genome-wide contact matrices from forward and reverse read .sam files (--binSize 500000; --restrictionSequence GATC; --danglingSequence GATC). HicSumMatrices was then used to create two merged .h5 genome-wide contact matrices: one containing two female samples and one containing four male samples. Afterwards, hicCorrectMatrix was run to give corrected genome-wide contact matrices (containing only one resolution/bin size) (--filterThreshold -1.5 3). From this point, multiple hicexplorer tools could be used. Given a .bw file for an active histone mark (H4K20me1), hicPCA was used to generate a Pearson’s correlation matrix and .bw eigenvectors/principle component files for each corrected matrix (--numberOfEigenvectors 1; --format bigwig; --histonMarkType active). Hicexplorer plotting commands were then used. HicPlotTADs can take .h5 corrected matrix files and .bw eigenvector files and provides figures with tracks showing the Hi-C matrix and eigenvector for a given region of the genome. HicPlotMatrix was used with the male .h5 corrected matrix file to generate plots of the sex chromosomes (--chromosomeOrder X1 Y1 X2 Y2 X3 Y3 X4 Y4 X5 Y5; --log1p).

### ChIP-seq peak calling

Macs2^10^ version 2.2.6 callpeak was used to call peaks from the .bam files of mapped ChIP-seq reads (-f BAM; -g 1.8e9; -B; -q 0.05). Histone peaks were called as either broad or narrow based on reference material available on the ENCODE project website (https://www.encodeproject.org/chip-seq/histone/).

### DNA methylation

Percentage methylation was presented in comparison with Hi-C and ChIP-seq data. Percentage methylation data originated from previous work^11^.

### Data and code availability

All sequencing data are available under BioProject PRJNA929280. The mass spectrometry proteomics data have been deposited to the ProteomeXchange Consortium via the PRIDE^1^ partner repository with the dataset identifier PXD040182. The scripts for the statistical models and the datasets necessary to run analyses included in this paper have been deposited in the public depository Git Hub, and are available at: < https://github.com/kango2/dcomp >

## Acknowledgements

Not applicable.

## Author Contributions

NCL drafted the manuscript and figures, performed data analysis, performed protein extraction on platypus heart and liver tissue samples. AMM performed data analysis. HRP performed data analysis and manuscript editing. SAW performed manuscript editing. BJH performed data analysis and manuscript editing. KLM performed manuscript editing. AML performed ChIP experiments. LKW performed experiments. ARR performed ChIP experiments as well as Hi-C experiments. SM was involved in editing of manuscript. MIR was involved in editing of manuscript. LSW was involved in editing of manuscript. FG was involved in editing of manuscript. JAMG was involved in editing of manuscript. ARH was involved in editing of manuscript and data analysis. PDW conceived and oversaw the project and the assembly of the manuscript.

## Additional information

Supplementary information containing protein summaries for platypus and opossum fibroblasts are available as excel spreadsheets. Files are named according to sample labelling described in ProteinPilot software workflow section of methods.

All other mass spectrometry data for platypus, chicken and mouse heart and liver samples, including raw files and protein summaries, are available at the ProteomeXchange Consortium via the PRIDE partner repository with the dataset identifier PXD040182.

## Extended data figure legends

**Extended Data Table 1.** Antibodies for ChIP-seq.

|  |  |  |
| --- | --- | --- |
| rabbit anti-H3K27ac | Diagenode | C15410174 |
| rabbit anti-H3K4me1 | Diagenode | C15410037 |
| rabbit anti-H3K4me3 | Merck Millipore | 07-473 |
| Rabbit anti-H4K20me1 | Abcam | ab9051 |

**Extended Data Table 2.** Reagents for HiC.

|  |  |  |
| --- | --- | --- |
| Biotin-14-dATP-50 nmol | Thermo Fisher Scientific | 19524016 |
| cOmplete, Mini, EDTA-free Protease Inhibitor Cocktail | SIGMA-ALDRICH | 4693159001 |
| DNA Pol. Large Fragm. (Klenow) | New England Biolabs | M0210L |
| Dynabeads MyOne Streptavidin T1-10 mL | Thermo Fisher Scientific | 65602 |
| NEBNext® Quick Ligation Reaction Buffer (5X) | New England Biolabs | B6058S |
| NEBNext® Ultra II Q5® Master Mix | New England Biolabs | M0544L |
| T4 DNA Ligase | New England Biolabs | M0202L |
| T4 DNA Polymerase | New England Biolabs | M0203L |
| T4 Polynucleotide Kinase NK | New England BioLabs | M0201 |
| NEBNext Multiplex Oligos for Illumina kit | New England Biolabs | E7500 |
| Quick Ligation™ Kit | New England Biolabs | M2200S |

**Extended Data Fig 1.**
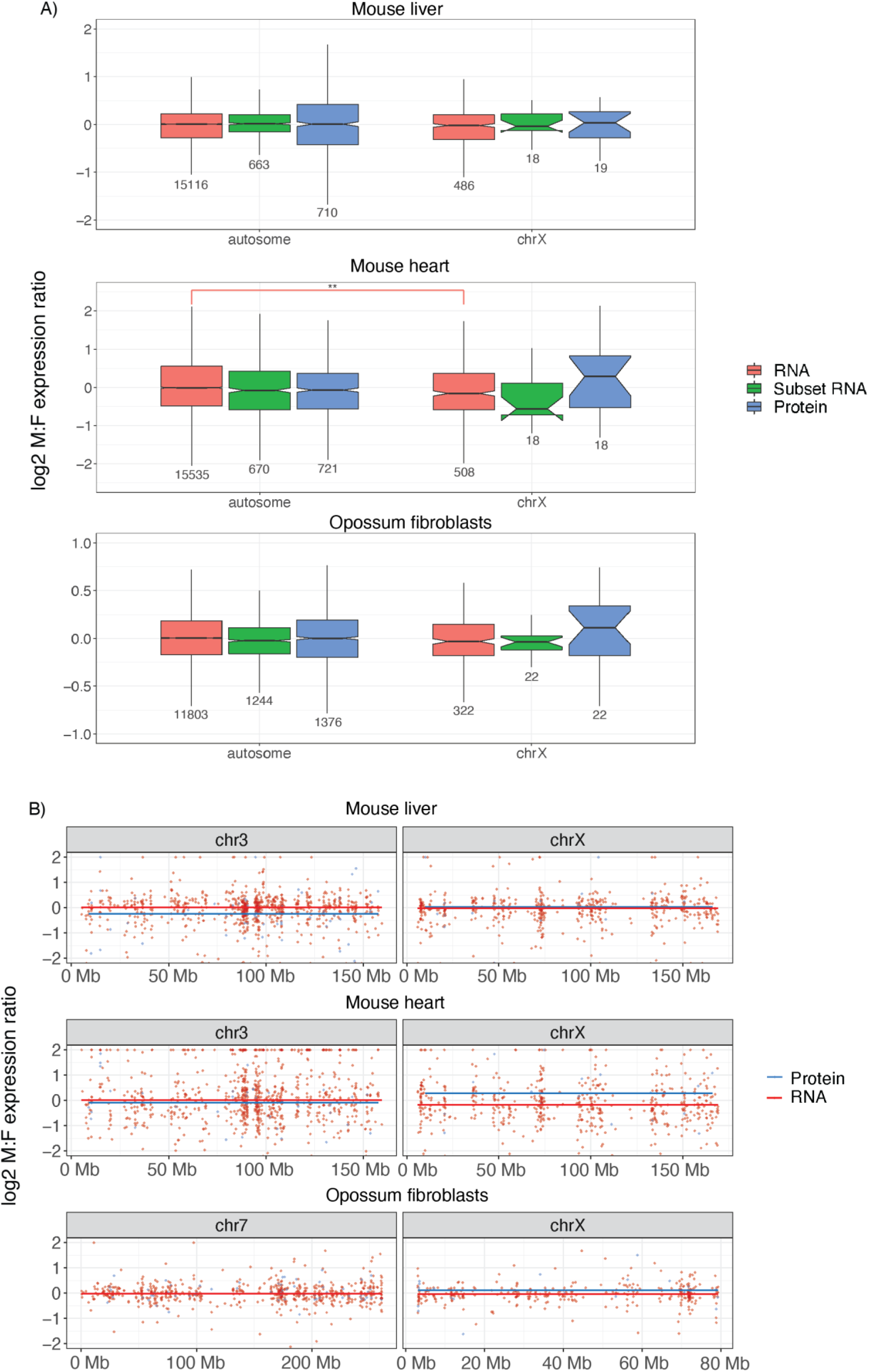
Male to female expression ratios of X-borne and autosomal genes in mouse and opossum. Median male to female (M:F) expression ratios (log2 scale) of autosomal and X borne genes in **A**) mouse heart and liver, and opossum fibroblasts. Ratios were calculated for all expressed mRNA (red), the subset of mRNAs sampled in the proteome (green), and proteins (blue). A ratio above zero is higher expression in males, whereas below zero is lower expression. Boxes represent the middle 50% of the data, and whiskers represent 1.5 times the interquartile range. Outliers are not plotted. Median is plotted inside the box, with the number of genes sampled below each boxplot. Mood’s median test was used to calculate if ratios for the autosomes and X were statistically different (** p < 0.001). **B**) Whole chromosome plots of M:F ratios for individual genes in the transcriptome (red) and proteome (blue) on the X and a representative autosome in mouse heart and liver, and opossum fibroblasts. Lines are the median M:F ratios for the whole chromosome.

**Extended Data Fig 2.**
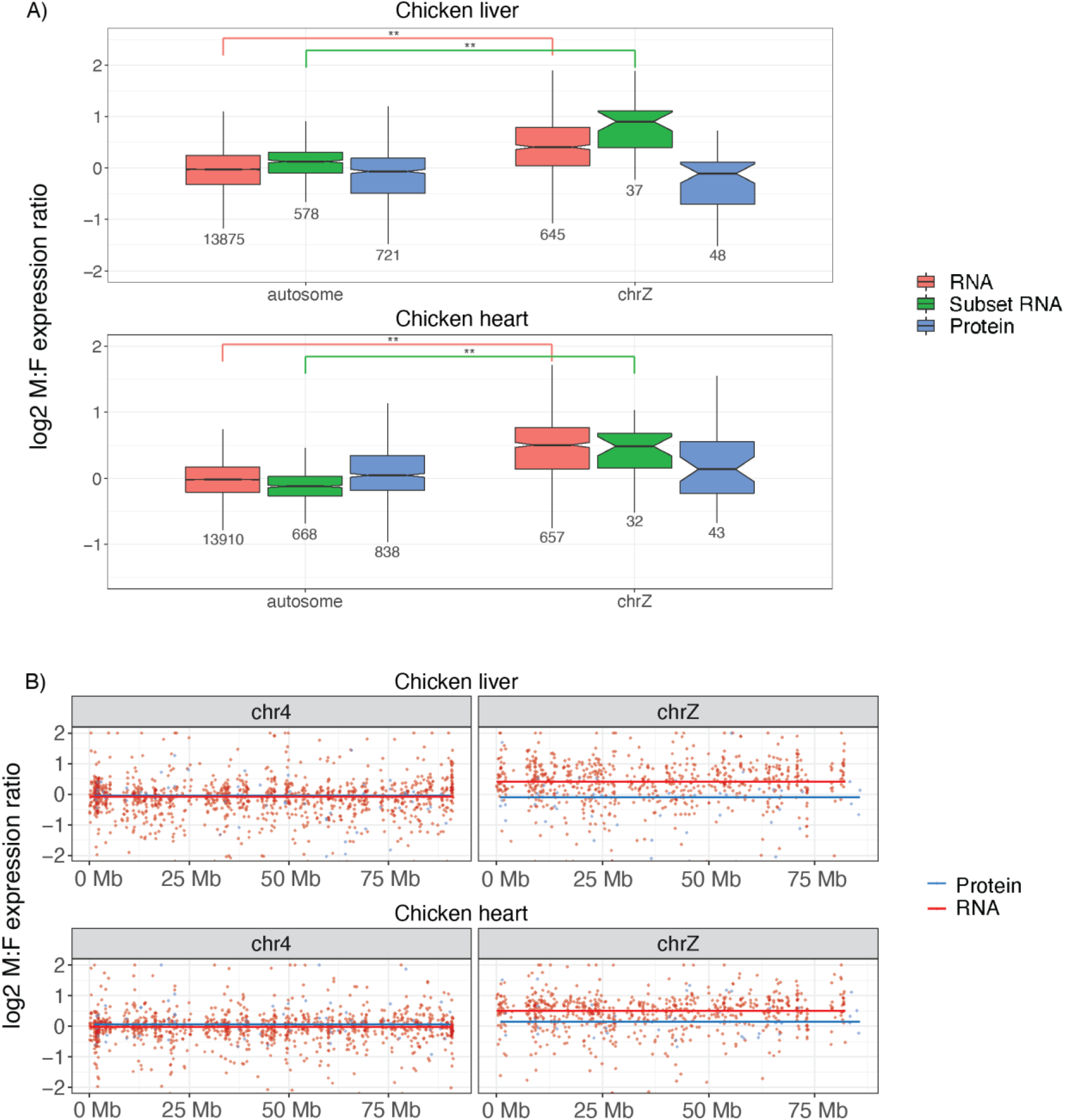
Male to female expression ratios of Z-borne and autosomal genes in chicken. Median male to female (M:F) expression ratios (log2 scale) of autosomal and X borne genes in **A**) chicken heart and liver. Ratios were calculated for all expressed mRNA (red), the subset of mRNAs sampled in the proteome (green), and proteins (blue). A ratio above zero is higher expression in males, whereas below zero is lower expression. Boxes represent the middle 50% of the data, and whiskers represent 1.5 times the interquartile range. Outliers are not plotted. Median is plotted inside the box, with the number of genes sampled below each boxplot. Mood’s median test was used to calculate if ratios for the autosomes and X were statistically different (** p < 0.001). **B**) Whole chromosome plots of M:F ratios for individual genes in the transcriptome (red) and proteome (blue) on the Z and a representative autosome in chicken heart and liver. Lines are the median M:F ratios for the whole chromosome.

**Extended Data Fig 3.**
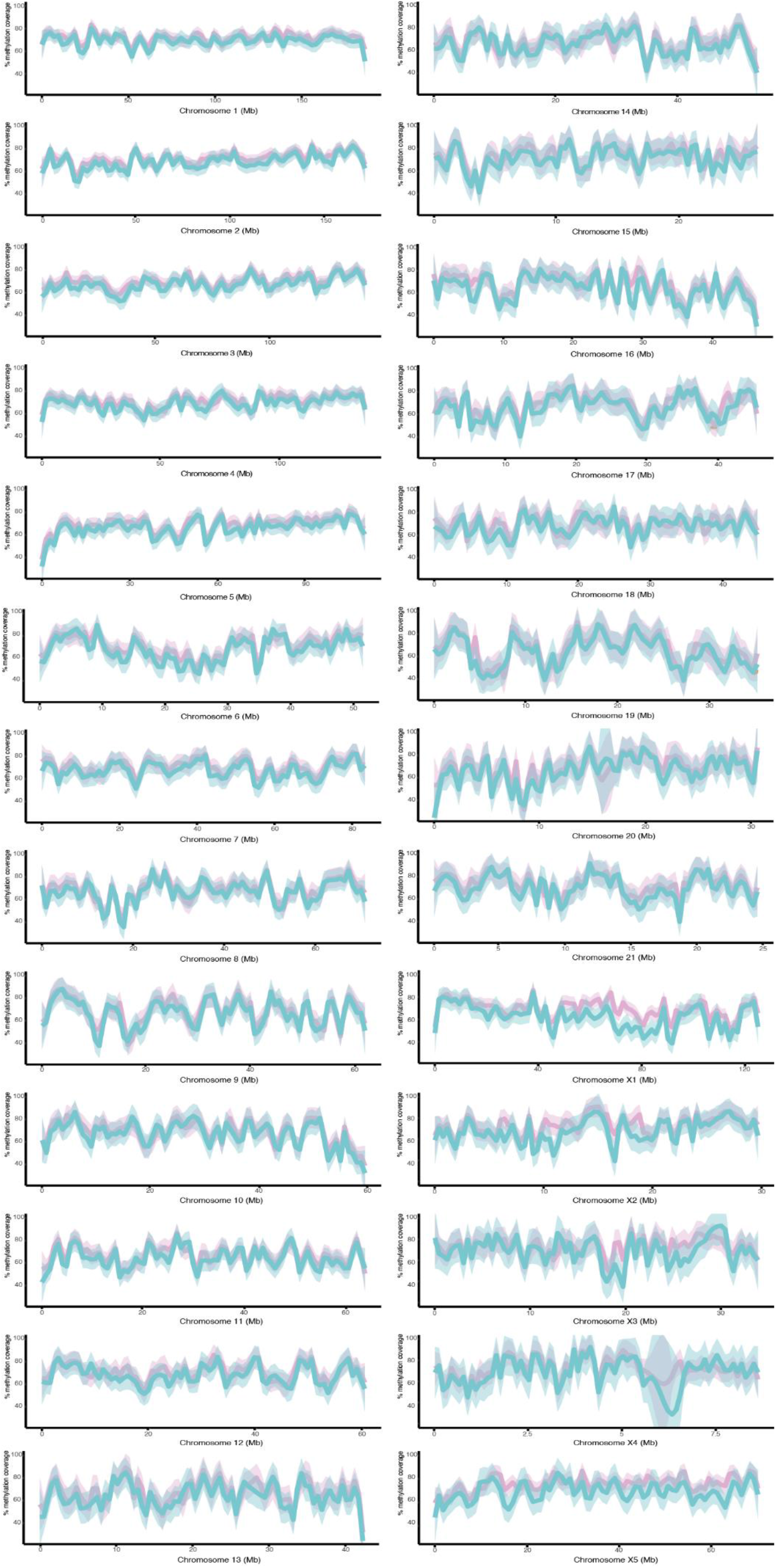
DNA methylation plots. Plots showing percent DNA methylation for all chromosomes in platypus, calculated in 50 kb non-overlapping tiles. Percent DNA methylation is represented as a smoothed line in pink for female and blue for male, with a shaded 95% confidence interval.

**Extended Data Fig 4.**
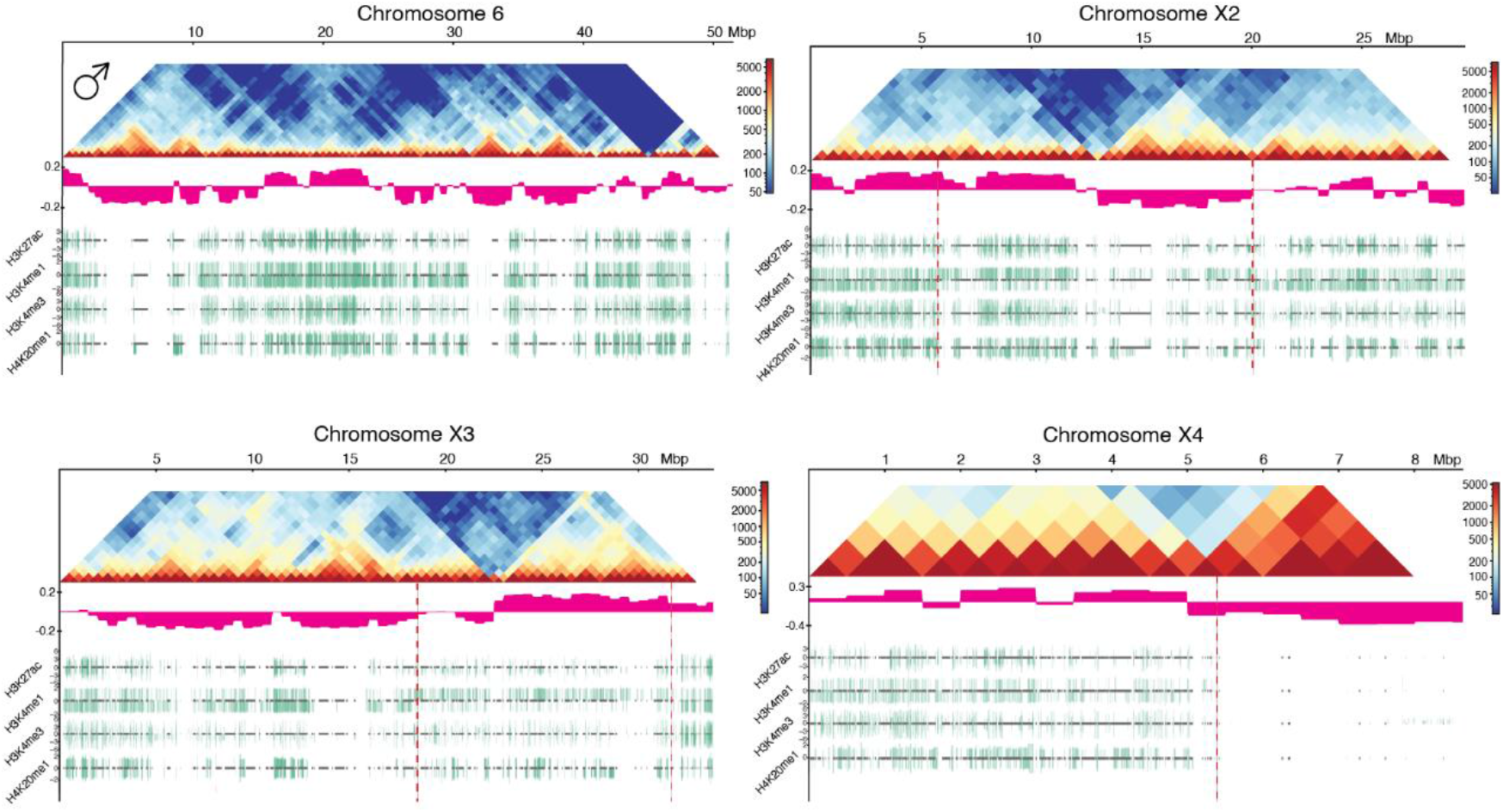
Epigenetic profile and 3D structure of chromosomes 6, X1, X2, X3 and X4 in platypus. Whole chromosome plots for chromosomes 6 (a representative autosome), X1, X2, X3 and X4 in platypus. The x-axis represents the full length of the chromosome, with size (in Mbp) at the top. Each plot shows a Hi-C contact matrix for male (top), generated using 500 kb bins. The Hi-C heatmap scale indicates number of contacts between bins. Below the Hi-C heatmaps are the eigenvectors in pink, which indicate A (positive) and B (negative) compartments. Below the male eigenvectors are ChIP-seq peak tracks in green for four active histone marks: H3K27ac, H3K4me1, H3K4me3 and H4K20me1. For each histone mark, male and female peaks are shown above and below the centre line respectively. ChIP peaks were filtered using a threshold q-value of 0.05 and peak height was displayed up to a fold change of 5. Gene locations are shown in the centre of the track as grey boxes. Lines are the median M:F ratios for the whole chromosome. On the X chromosome plots, vertical dashed red lines indicate PAR boundaries. PAR boundaries were identified using the Hi-C data.

**Extended Data Fig 5.**
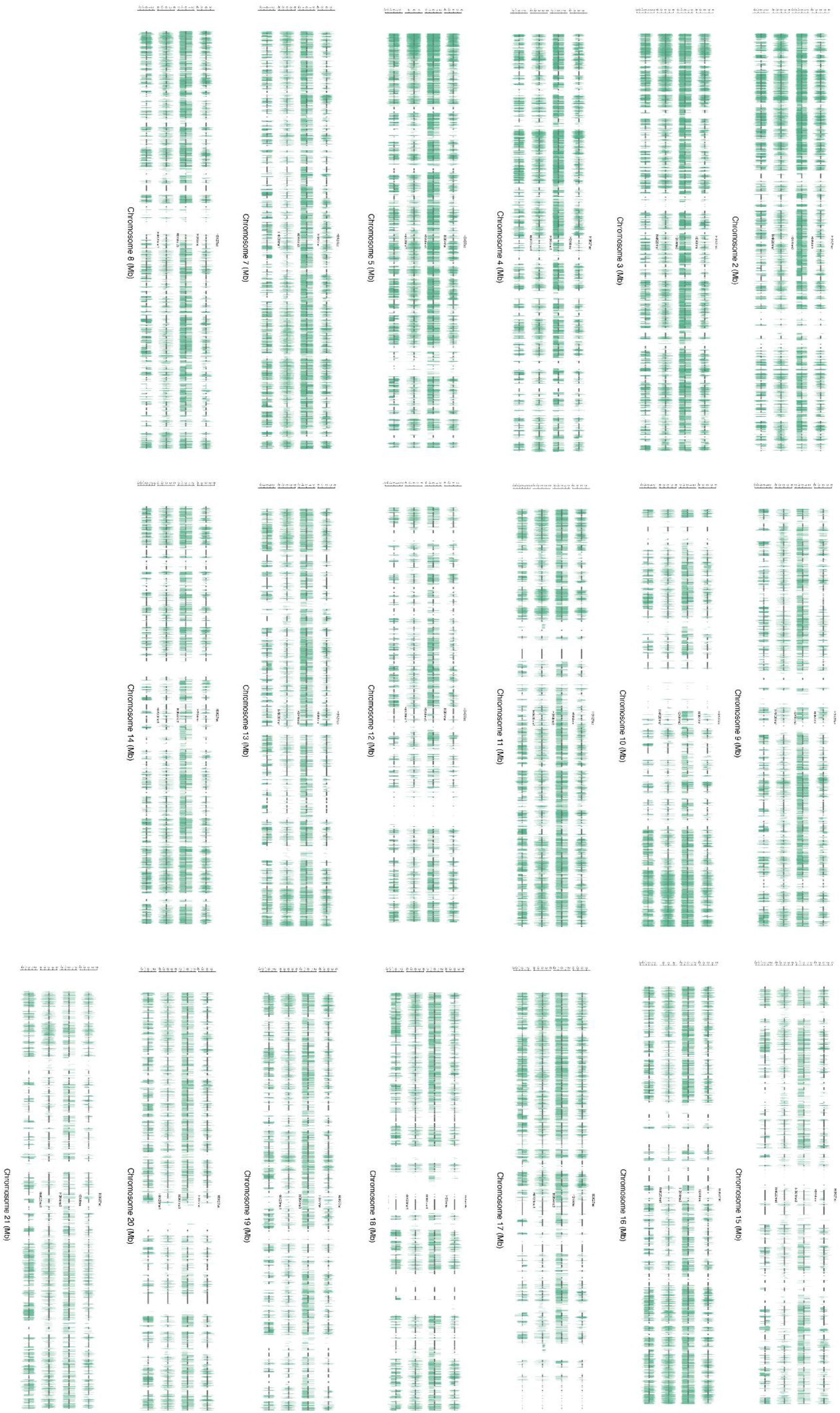
ChIP-seq peaks. ChIP-seq peak tracks for chromosomes 2-21 (excluding chromosome 6, see Figure S2) in platypus. For each chromosome, four active histone marks are shown (from top to bottom): H3K27ac, H3K4me1, H3K4me3 and H4K20me1. For each histone mark, male and female peaks are shown above and below the centre line respectively.

